# DNA barcodes and insights into the phylogenetic relationships of butterflies of the genus *Eurema* (Pieridae) from Uttarakhand, India

**DOI:** 10.1101/242263

**Authors:** Ankita Rajpoot, Ved Prakash Kumar, Archana Bahuguna

## Abstract

In DNA barcoding, mitochondrial gene cytochrome *c* oxidase I recommended as a tool for the rapid identification and discovery of species. Genus *Eurema*, Family Pieridae is a highly diverse and distributed along wide geographic ranges in the world as well as India including approximately 70 species throughout the world. The present study is preliminary approach, we included n=12 specimen (3 samples per species) of four different *Eurema* species, listed in IUCN as Least concern species, were collected from Uttarakhand (India), to give the DNA barcodes and examine patterns of gene evolution through molecular phylogenetics with publicly available sequences of other 17 *Eurema* species present in different countries.

The generated (n=12) COI sequences compared with the sequences of the con-specifics submitted from different geographic regions, all four species were correctly identified. The obtained COI barcodes clearly showed the intraspecific and interspecific distance among four *Eurema* species by using a K2P technique. In spite NJ, clustering analysis also successfully discriminated all four species.

In phylogenetic topology, included 21 *Eurema* species in the Bayesian and Maximum Likelihood analysis recovered all species in two monophyletic clades with strong support and resolved taxonomic position within species groups. Although higher-level relationships among other *Eurema* species groups require additional study.

---

DNA barcoding (http://www.barcodinglife.org) has gained so much popularity during the last decade in documenting the global biodiversity (Hebert et al. 2003ab, Ebach and Holdrege 2005, Abdo and Golding 2007, Meusnier et al. 2008, Monaghan et al. 2009, Hajibabaei et al. 2011, Haye et al. 2012, Leray and Knowlton 2015). Cytochrome *c* oxidase subunit I (COI) is a widely used standard mitochondrial gene for DNA barcoding to facilitate identification of biological specimens (Hebert et al. 2003). DNA Barcoding is also using as a tool for discrimination between cryptic species and to discover new species based on nucleotide sequence divergence. Availability of large amount of sequences in public databases will indeed speed up species description and identification for taxonomist (Blaxter 2004, Ratnasingham and Hebert 2007). Overall the goal of DNA barcoding is explicitly to aid species identification, it has frequently been used for phylogenetic inference at multiple taxonomic levels (Tautz et al. 2003, Savolainen et al. 2005) prompting many scientist to contemplate the phylogenetic value of DNA barcode datasets (Tautz et al. 2003, Savolainen et al. 2005, Hajibabaei et al. 2007, Wahlberg and Wheat 2008).

Uttarakhand lies in Central Himalaya at 77°45’ East longitudes to 81° North longitudes, elevation ranges from 300 m to 7000 meters above sea level covering a geographical area of 53,485 sq km, has a great diversity of flora and fauna constitutes 65% of the total area of the state (Sundriyal and Sharma 2016, Uttarakhand Annual Plan 2011–12). The rich faunal and floral diversity of the state comprises 4907 faunal species (include 3948 invertebrate and 959 vertebrate species) and 5096 Floral species (included Angiosperms and Gymnosperms) (Arora and Kumar 1995, Tak and Sati 2010, Envis Newsletter 2013). High floral diversity indicates high butterfly diversity in state and butterflies is an important group of ‘model' organisms used, for centuries, to investigate many areas of biological research, ecology, evolution, population genetics and developmental biology (Boggs et al. 2003).

Lepidoptera, as the second largest order of insects with more than 157,000 species, are of particular interest in systematic research. Among them, India is home to about 1800 species and subspecies of butterflies (Kunte et al. 2017) which is about 8.74% of total butterfly species in the world and constitute 65% of total Indian fauna. Uttarakhand harbours 407 species of butterflies (Singh and Sondhi 2016). Due to their beautiful colours and high diversity, butterflies are well studied through morphologically, systematically, and ecological perspective in India as well as Uttarakhand (Kunte 2000, Khan et al. 2004, Kumar 2008, Jain and Jain 2012, Bhardwaj et al. 2012, 2013, Kumaraswamy and Kunte 2013, Kunte et al. 2015). Family Pieridae is highly diverse and distributed along wide geographic ranges. Unfortunately, there are few molecular studies has been reported on butterflies from India (Ragupathy et al. 2009, Gaikwad et al. 2011).

Genus *Eurema* (Hübner 1819) is highly diverse genus of family Pieridae, distributed along wide geographic ranges in the world as well as India including approximately 70 species throughout the world (*Eurema*, page). *Eurema* butterflies are comparatively small butterflies with lemon yellow wings, bordered with black, especially on the upper side of both wings, commonly called grass yellow butterflies. The undersides of the wings, which are usually light in colour than the upper side, are marked with several black or faint brown markings (Osamu Yata 1991, Jeratthitikul et al. 2009). There are number of molecular studies available on different family of butterflies including DNA barcoding, molecular phylogeny and phylogeography aspect (Scott 1985, Brower and DeSalle 1998, Zimmermann et al. 2000, Xue 2009, Xiangqun et al. 2015, Martin et al. 2017). Although few molecular studies available on family Pieridae (Braby et al. 2006, Solovyev et al. 2015).

In India there are eight species reported under the genus *Eurema* such as *Eurema brigitta* Cramer, 1780 (Small grass yellow); *E. laeta* Boisduval, 1836 (Spotless grass yellow); *E. lacteola* Distant, 1886 (Scare grass yellow); *E. andersonii*, Moore (One-spot grass yellow); *E. hecabe* Linnaeus, 1758 (Common grass yellow); *E. blanda* Boisduval, 1836 (Three-spot grass yellow), *E. nilgiriensis* Menon, 1987 (Nilgiri grass yellow) and *E. simulatrix* Semper, 1891 (Scarce Changeable Grass Yellow) (Kunte 2017). Out of these, five species (*E. brigitta, E. laeta, E. andersonii, E. hecabe*, and *E. blanda*) are reported from Uttarakhand (Smetacek 2013, Singh and Sondhi 2016). Among, these two species (*E. brigitta and E. andersonii*) listed as least concern in IUCN (Larsen 2011, Muller and Tennent 2011). In present study, we included five *Eurema* species (4 from current study and 1 from Genbank) from Uttarakhand to develop an authentic genetics reference library based on DNA barcoding approach. Furthermore, we conducted phylogenetic analysis to clarify the relationships between the other species of genus *Eurema*, which is globally distributed through using the sequences available in GenBank.

## MATERIALS AND METHODS

### Sampling for Genetic analysis

N=12 samples (3 samples per species) from each four species of genus *Eurema*, were collected from various localities in the Mussoorie and Dehradun, Uttarakhand, India (Fig. 1) Additionally, we also used another (n=78) sequences of 17 *Eurema* species including a Uttarakhand specie (*E. andersoni*) from NCBI Genbank for the phylogenetic analysis (Table 1). Butterflies were collected by using the sweeping net and dead or damaged animals then placed in a separate glassine envelope stored at −20°C until further use. One or more legs removed, kept in ethanol for DNA isolation. For initial identification, we evaluate the wing marking patterns of the all species found in Uttarakhand except *E. andersonii* (A rare species, Reported in 2003, 2004 and 2015 (Singh and Bhandari 2003, Singh and Sondhi 2016). *It* earlier reported by Mackinnon and de Nicéville (1899) from Mussoorie, Uttarakhand.

**Table 1.**
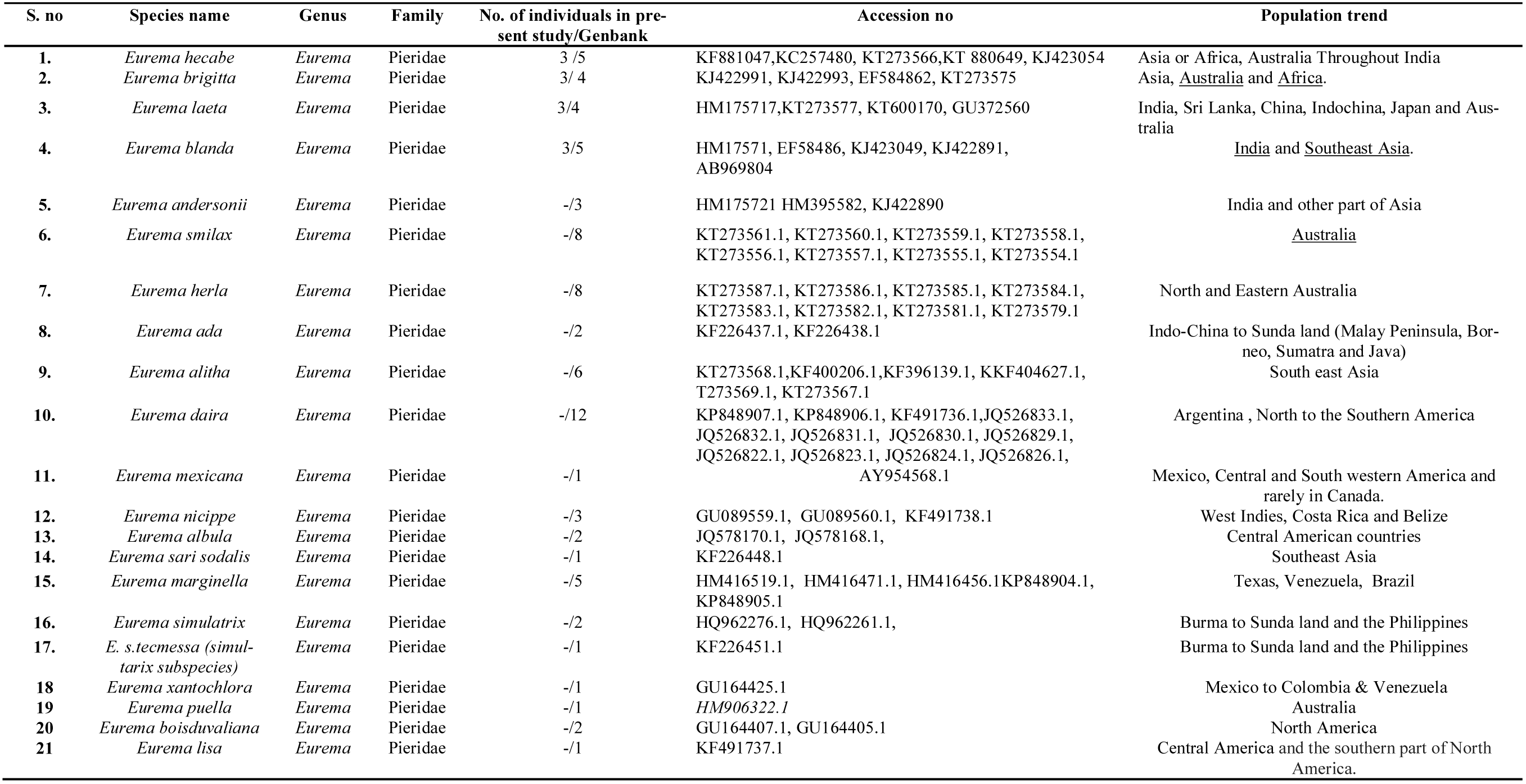
List of *Eurema* species, number of individuals, accession numbers and population trend of each used in the present study

**Figure 1.**
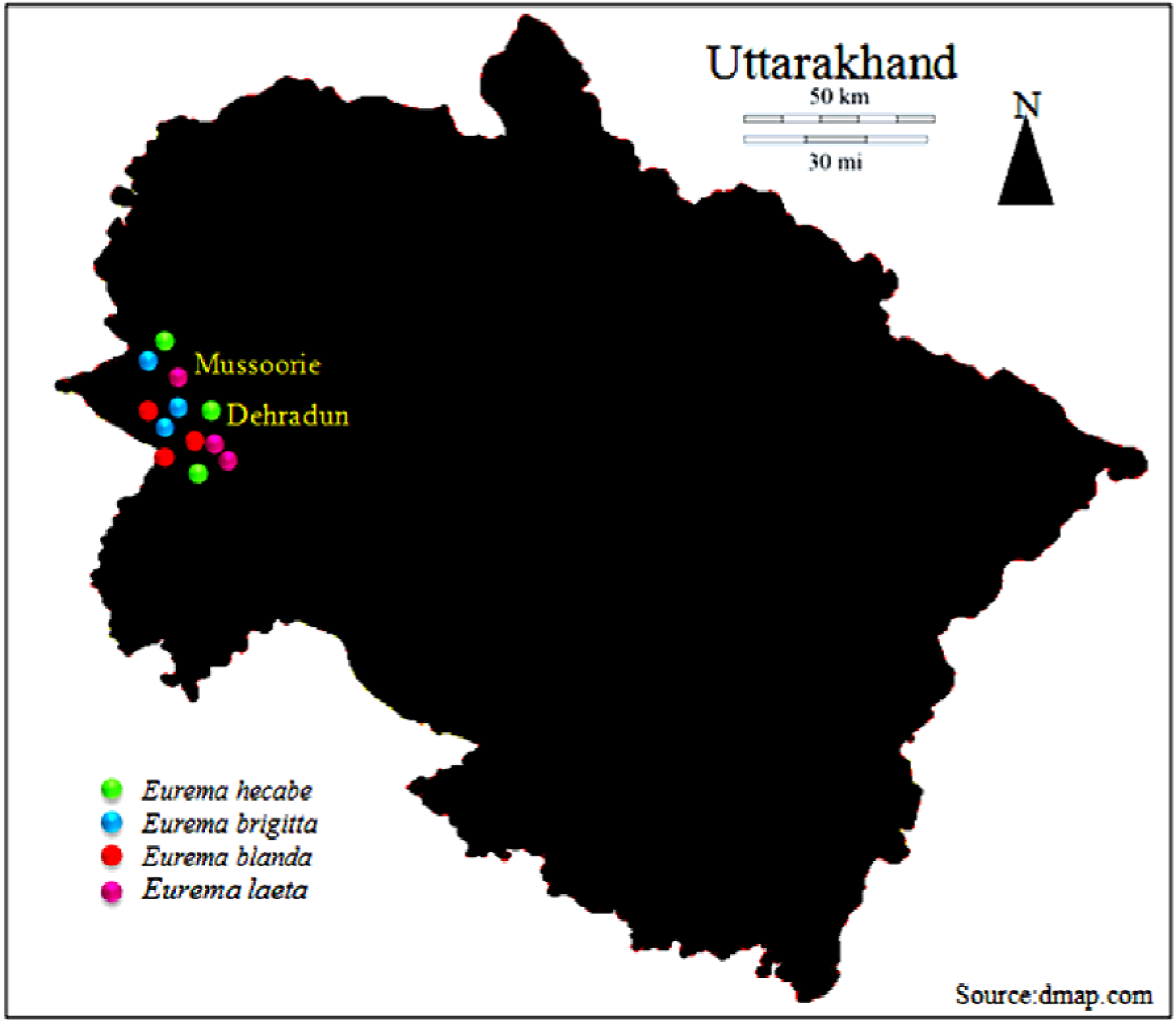
Sampling location of four *Eurema* species included in present study from Uttarakhand.

### Laboratory procedure

The total genomic DNA was extracted from the leg samples (n=12) using Qiagen DNeasy Tissue Kit (QIAGEN, Germany) protocol; a partial fragment of COI (Folmer et al. 1994) was amplified in an Eppendorf Master Thermocycler. Polymerase chain reaction (PCR) master mix preparation and PCR cycling condition performed according to the previously published article (Rajpoot et al. 2017). To check the contamination of the DNA, a negative control was set up with a PCR master mix. Sanger sequencing performed by a commercial using the same primers. All obtained sequences were good for both the reverse and forward primers. Chromatograms in both directions compared using Codon Code Aligner 3.9 (Codon Code Corporation) and automatic base calls checked along the sequence, both where the two sequences were in disagreement and elsewhere.

### Data analysis

The sequences obtained were validated and cleaned using Cromas 2.6.4 (http://www.technelysium.com.au), multiple sequence alignment with high accuracy and high throughput sequences of equal length were generated after multiple sequence alignment (MSA) using MUSCLE (Edgar 2004), sequences characteristic were performed manually software MEGA 7 (Kumar et al. 2016). FASTA format of these species sequences conformed by BLAST search at NCBI. Moreover, recheck the BLAST result, neighbor-joining (NJ) (Saitou and Nei 1987) with bootstrap test (1000 replicates) (Felsenstein 1985) using to see the closet taxa. The Kimura 2-parameter (K2P) model (Kimura 1980) of base substitution was used to calculate pairwise genetic distance in MEGA 7 software (Kumar et al. 2016). Additionally, to check the performance of DNA barcoding, we downloaded sequences of same species from Genbank originated from different geographical areas (Table 1). The intra and interspecies nucleotide divergence and haplotypes variation calculated using MEGA 7 (Kumar et al. 2016).

Phylogenetic analysis was performed using MrBayes v3.1 (Ronquist et al. 2011) to establish relationships based on Bayesian Inferences (BI), while a Maximum Likelihood (ML) approach was implemented in PhyML v3.1 (Guindon et al. 2010). The Akaike Information Criterion (AIC) in jModelTest v2.1.3 (Darriba et al. 2012) employed to determine the best-fit model of sequence evolution. The same model selected for BI and ML analysis as it proved to be the best-fit model calculated for each analysis. Nodal support for the ML analyses carried out with 1000 bootstrap replications. The BI calculations were run over two million generations, after which 25% of the trees were discarded as burn-in. Tracer v1.5 (Drummond and Rambaut, 2007) software was used to assess trace files generated by MrBayes v3.1, in order to assess whether mixing was achieved and to choose a suitable percentage burn-in. Moreover to find out the more accurate tree topology *Pontia edusa* (*KY846094.1*) was used as an outgroup (Xiangqun et al. 2015).

## RESULT AND DISCUSSION

### Morphological identification

The collected 12 specimens of genus *Eurema* species first confirmed based on the morphological characteristics. Identification done by observing wing shape and colour pattern described in available keys/identification guides (Antram 1924, Peile 1937, Gunthilagaraj et al. 1998, Kunte 2000, Rangnekar 2007).

### Analysis of molecular dataset

Isolation, PCR amplification and sequencing of DNA from leg of butterflies yielded good quality sequences. We generated DNA barcode sequences of 12 specimens representing four distinct, morphologically identifiable species, belonging to a single genus *Eurema*, the obtained length of these COI sequences were 565 bp. All sequences generated in this study submitted in GenBank (sequence submitted but waiting for Accession number) for future uses.

Most of our sequences 98%-100% matched with the COI sequences of respective species, which already deposited in NCBI GenBank databases (Table 2). The obtained sequences of 565 bp (COI gene) of 12 specimen contained 409 Conserved regions, 92 Variable sites, 75 Parsimony informative sites, 16 Singleton sites and 8 haplotypes (Table 2). The maximum likelihood estimate of transition /transversion bias (R) were 1.60 and nucleotide composition were A 38.6, T 30.6, C 13.6, G 17.1 (Fig. 2). Furthermore, the overall haplotypes (hd) and nucleotide diversity (π) was 0.923 and 0.08807 respectively, while, the overall sequences divergence (k) among all sequences was 0.93% and overall nucleotide difference between four species was 43.765 (Table 2).

**Table 2.**
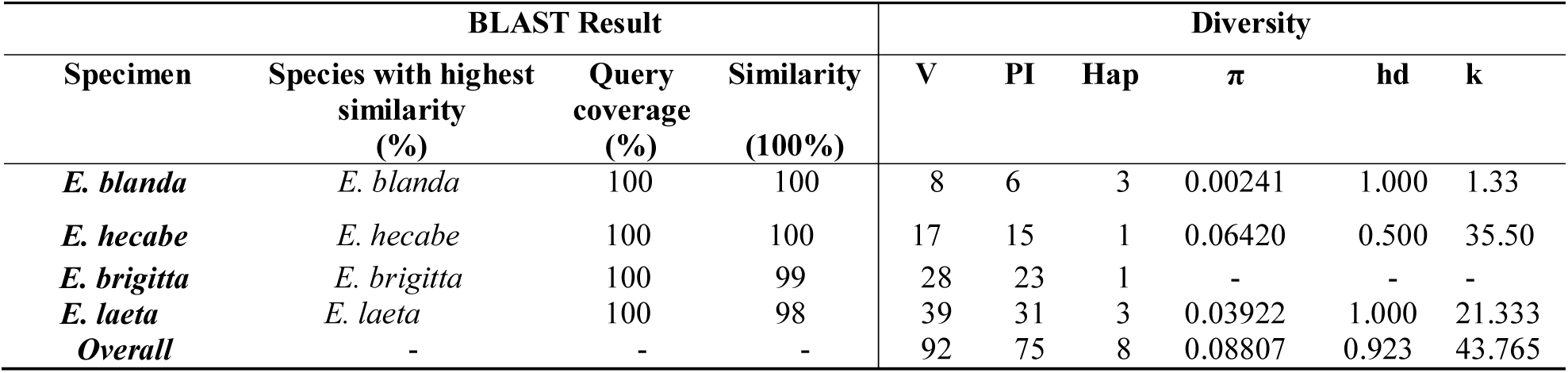
Species cross verification and diversity estimates of four *Eurema* species of Uttarakhand. Variable sites (V); Parsimony Informative sites (PI); Haplotypes (Hap); Nucleotide diversity (π); Haplotype diversity (hd); overall sequences divergence (k)

| BLAST Result |  |  |  | Diversity |  |  |  |  |  |
| --- | --- | --- | --- | --- | --- | --- | --- | --- | --- |
| Specimen | Species with highest similarity (%) | Query coverage (%) | Similarity (100%) | V | PI | Hap | $\pi$ | hd | k |
| *E. blanda* | *E. blanda* | 100 | 100 | 8 | 6 | 3 | 0.00241 | 1.000 | 1.33 |
| *E. hecabe* | *E. hecabe* | 100 | 100 | 17 | 15 | 1 | 0.06420 | 0.500 | 35.50 |
| *E. brigitta* | *E. brigitta* | 100 | 99 | 28 | 23 | 1 | - | - | - |
| *E. laeta* | *E. laeta* | 100 | 98 | 39 | 31 | 3 | 0.03922 | 1.000 | 21.333 |
| **Overall** | - | - | - | 92 | 75 | 8 | 0.08807 | 0.923 | 43.765 |

**Figure 2:**
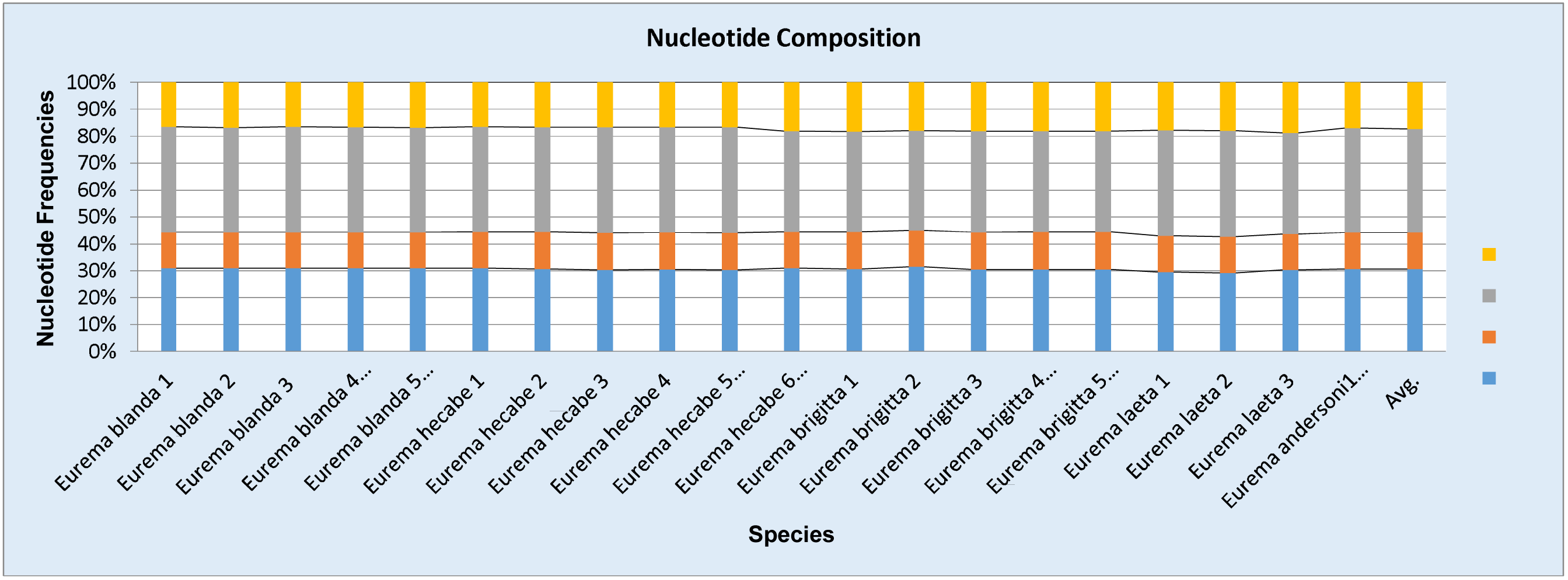
Observed nucleotide frequencies among 5 *Eurema* species (from Uttarakhand) in mitochondrial COI gene

### Determination of Intraspecific and Interspecific K2P distances

The obtained COI sequences clearly showed the intraspecific and interspecific distance among four *Eurema* species by using a K2P technique. The interspecific variation in *Eurema*, mean pairwise sequences divergences (d) of COI ranged from 0.061% (*E. blanda/E. hecabe*) to 0.131% (*E. hecabe/E. leata*) with an average genetic distance of 0.84%. Although, *d* value 0.118% and 0.112% occur in *E. blanda/E. brigitta* and *E. blanda/E. leata* respectively, *d* value 0.114% and 0.119% occur in *E. hecabe/E. brigitta* and *E. brigitta/E. leata* correspondingly. In addition, observed overall interspecific mean diversity among four species was 0.06% (Table 3).

**Table 3.**
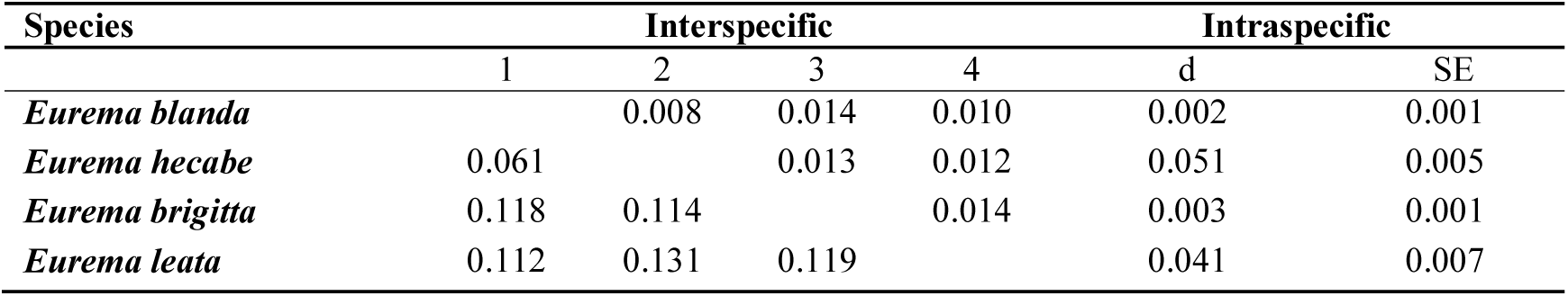
Interspecific evolutionary divergence between four *Eurema* species of Uttarakhand. The number of base substitutions per site from averaging over all sequence pairs between groups are shown. Standard error estimate(s) are shown above the diagonal and were obtained by a bootstrap procedure (1000 replicates). Analysis were conducted using the Kimura 2-parameter model.

| Species | Interspecific |  |  |  | Intraspecific |  |
| --- | --- | --- | --- | --- | --- | --- |
|  | 1 | 2 | 3 | 4 | D | SE |
| *Eurema blanda* |  | 0.008 | 0.014 | 0.010 | 0.002 | 0.001 |
| *Eurema hecabe* | 0.061 |  | 0.013 | 0.012 | 0.051 | 0.005 |
| *Eurema brigitta* | 0.118 | 0.114 |  | 0.014 | 0.003 | 0.001 |
| *Eurema leata* | 0.112 | 0.131 | 0.119 |  | 0.041 | 0.007 |

The intraspecific variation, *d* value ranged from 0.002% (*E. blanda*) to 0.051% (*E. hecabe*) with an average genetic distance 024%. However, d value 0.003% and 0.041% occurred in *E. brigitta* and *E. leata* respectively. Moreover, observed overall intraspecific mean diversity was 0.09% (Table 3).

To reconfirmed, the interspecific and intraspecific evolutionary divergence within Uttarakhand *Eurema* species, K2P based NJ tree topology showed three well supported monophyletic clades, where *E. blanda* and *E. hecabe* clustered together in the first clade with 65% to 100% bootstrap value, *E. andersonii* present in second clade with 95% bootstrap value from clade first, while *E. brigitta* and *E. leata* clustered together in third clade with 64% to 100% bootstrap value. Overall, tree topology supported the interspecific evolutionary divergences result where all five species showed barcoding variation (Fig. 3). Moreover, variation of species-specific sites between five species given in Table 5.

**Figure 3:**
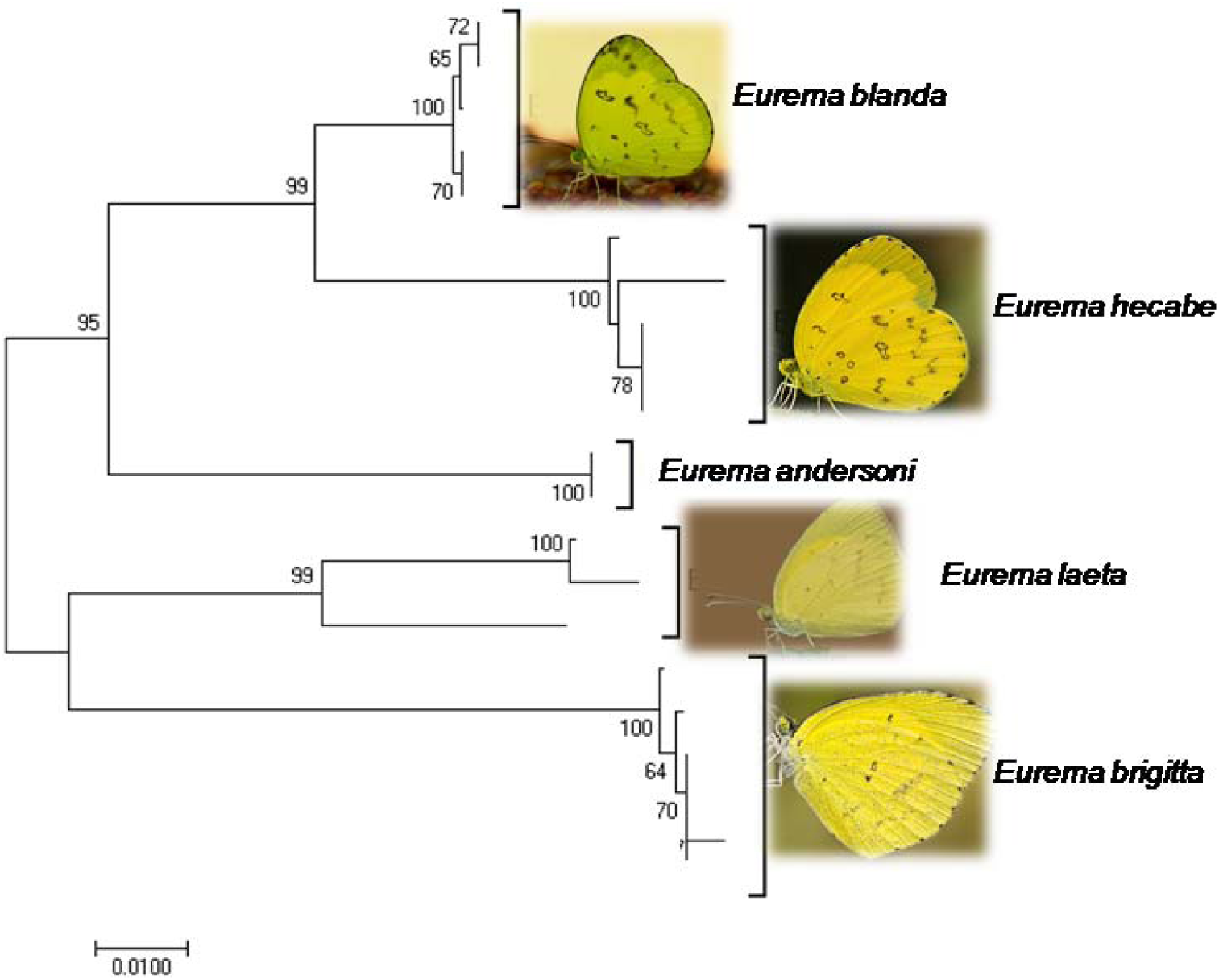
The COI sequences based Neighbor-Joining (NJ) tree among 5 *Eurema* species Uttarakahnd. The percentage of replicate trees in which the associated taxa clustered together in the bootstrap test (1000 replicates) shown next to the branches. The evolutionary distances were computed using the Kimura 2-parameter method and are in the units of the number of base substitutions per site.

### Evolutionary divergence and phylogenetic relationship among twenty-one Eurema species

Calculation of mean pairwise sequences divergences (d) between different *Eurema* species showed that they ranged from 0.1% (*E. l .lacteola/E. hecabe/E. ada*) to 0.15% (*E. brigitta/E. lisa; E. nicippe/E. s. techmessa; E. s. techmessa/E. boisduvaliana/ E. lisa*) with an average genetic distance of 0.10% (Table 4).

**Table 4.**
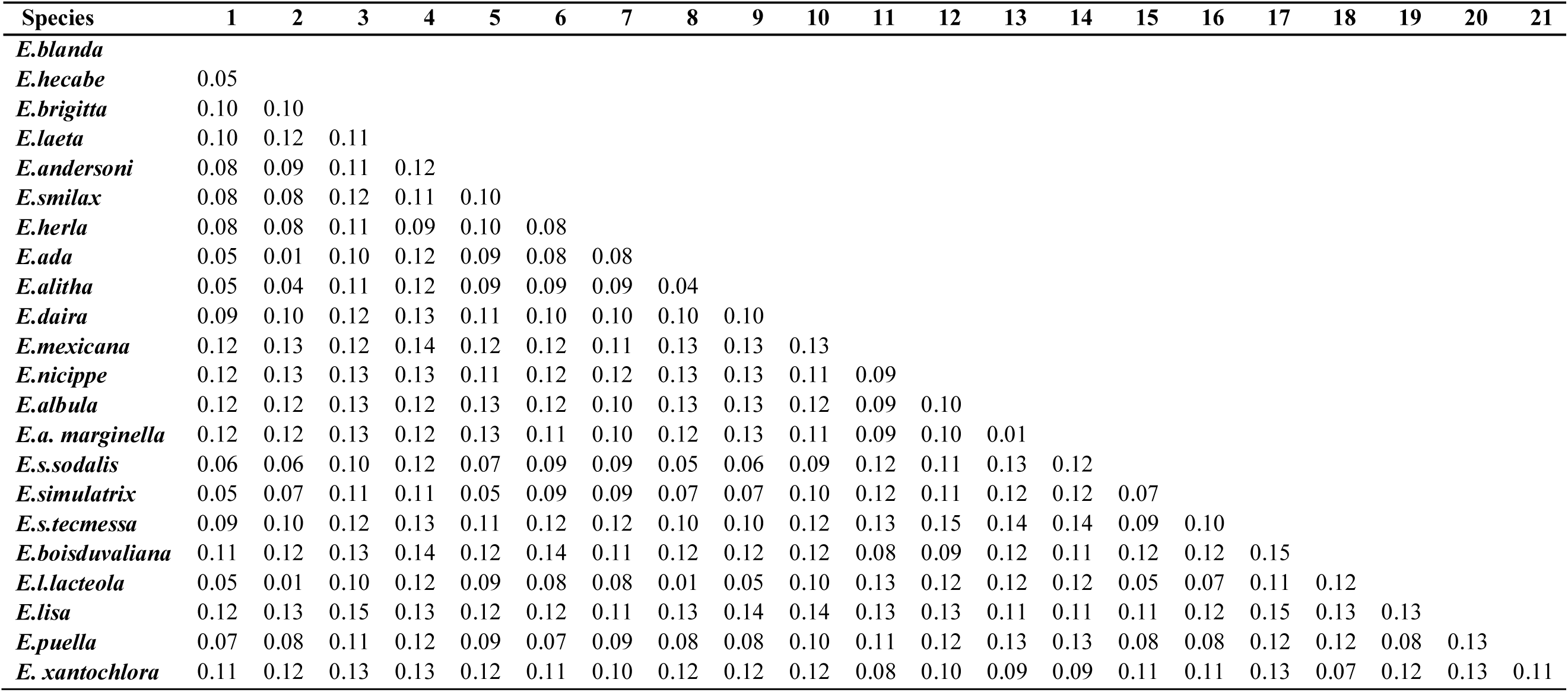
Estimates of evolutionary divergence over sequence pairs between 21 *Eurema* species. Maximum and minimum divergences highlighted in table 1

| Species | 1 | 2 | 3 | 4 | 5 | 6 | 7 | 8 | 9 | 10 | 11 | 12 | 13 | 14 | 15 | 16 | 17 | 18 | 19 | 20 | 21 |
| --- | --- | --- | --- | --- | --- | --- | --- | --- | --- | --- | --- | --- | --- | --- | --- | --- | --- | --- | --- | --- | --- |
| *E.blanda* |  |  |  |  |  |  |  |  |  |  |  |  |  |  |  |  |  |  |  |  |  |
| *E.hecabe* | 0.05 |  |  |  |  |  |  |  |  |  |  |  |  |  |  |  |  |  |  |  |  |
| *E.brigitta* | 0.10 | 0.10 |  |  |  |  |  |  |  |  |  |  |  |  |  |  |  |  |  |  |  |
| *E.laeta* | 0.10 | 0.12 | 0.11 |  |  |  |  |  |  |  |  |  |  |  |  |  |  |  |  |  |  |
| *E.andersoni* | 0.08 | 0.09 | 0.11 | 0.12 |  |  |  |  |  |  |  |  |  |  |  |  |  |  |  |  |  |
| *E.smilax* | 0.08 | 0.08 | 0.12 | 0.11 | 0.10 |  |  |  |  |  |  |  |  |  |  |  |  |  |  |  |  |
| *E.herla* | 0.08 | 0.08 | 0.11 | 0.09 | 0.10 | 0.08 |  |  |  |  |  |  |  |  |  |  |  |  |  |  |  |
| *E.ada* | 0.05 | 0.01 | 0.10 | 0.12 | 0.09 | 0.08 | 0.08 |  |  |  |  |  |  |  |  |  |  |  |  |  |  |
| *E.alitha* | 0.05 | 0.04 | 0.11 | 0.12 | 0.09 | 0.09 | 0.09 | 0.04 |  |  |  |  |  |  |  |  |  |  |  |  |  |
| *E.daira* | 0.09 | 0.10 | 0.12 | 0.13 | 0.11 | 0.10 | 0.10 | 0.10 | 0.10 |  |  |  |  |  |  |  |  |  |  |  |  |
| *E.mexicana* | 0.12 | 0.13 | 0.12 | 0.14 | 0.12 | 0.12 | 0.11 | 0.13 | 0.13 | 0.13 |  |  |  |  |  |  |  |  |  |  |  |
| *E.nicippe* | 0.12 | 0.13 | 0.13 | 0.13 | 0.11 | 0.12 | 0.12 | 0.13 | 0.13 | 0.11 | 0.09 |  |  |  |  |  |  |  |  |  |  |
| *E.albula* | 0.12 | 0.12 | 0.13 | 0.12 | 0.13 | 0.12 | 0.10 | 0.13 | 0.13 | 0.12 | 0.09 | 0.10 |  |  |  |  |  |  |  |  |  |
| *E.a. marginella* | 0.12 | 0.12 | 0.13 | 0.12 | 0.13 | 0.11 | 0.10 | 0.12 | 0.13 | 0.11 | 0.09 | 0.10 | 0.01 |  |  |  |  |  |  |  |  |
| *E.s.sodalis* | 0.06 | 0.06 | 0.10 | 0.12 | 0.07 | 0.09 | 0.09 | 0.05 | 0.06 | 0.09 | 0.12 | 0.11 | 0.13 | 0.12 |  |  |  |  |  |  |  |
| *E.simulatrix* | 0.05 | 0.07 | 0.11 | 0.11 | 0.05 | 0.09 | 0.09 | 0.07 | 0.07 | 0.10 | 0.12 | 0.11 | 0.12 | 0.12 | 0.07 |  |  |  |  |  |  |
| *E.s.tecmessa* | 0.09 | 0.10 | 0.12 | 0.13 | 0.11 | 0.12 | 0.12 | 0.10 | 0.10 | 0.12 | 0.13 | 0.15 | 0.14 | 0.14 | 0.09 | 0.10 |  |  |  |  |  |
| *E.boisduvaliana* | 0.11 | 0.12 | 0.13 | 0.14 | 0.12 | 0.14 | 0.11 | 0.12 | 0.12 | 0.12 | 0.08 | 0.09 | 0.12 | 0.11 | 0.12 | 0.12 | 0.15 |  |  |  |  |
| *E.l.lacteola* | 0.05 | 0.01 | 0.10 | 0.12 | 0.09 | 0.08 | 0.08 | 0.01 | 0.05 | 0.10 | 0.13 | 0.12 | 0.12 | 0.12 | 0.05 | 0.07 | 0.11 | 0.12 |  |  |  |
| *E.lisa* | 0.12 | 0.13 | 0.15 | 0.13 | 0.12 | 0.12 | 0.11 | 0.13 | 0.14 | 0.14 | 0.13 | 0.13 | 0.11 | 0.11 | 0.11 | 0.12 | 0.15 | 0.13 | 0.13 |  |  |
| *E.puella* | 0.07 | 0.08 | 0.11 | 0.12 | 0.09 | 0.07 | 0.09 | 0.08 | 0.08 | 0.10 | 0.11 | 0.12 | 0.13 | 0.13 | 0.08 | 0.08 | 0.12 | 0.12 | 0.08 | 0.13 |  |
| *E. xantochlora* | 0.11 | 0.12 | 0.13 | 0.13 | 0.12 | 0.11 | 0.10 | 0.12 | 0.12 | 0.12 | 0.08 | 0.10 | 0.09 | 0.09 | 0.11 | 0.11 | 0.13 | 0.07 | 0.12 | 0.13 | 0.11 |

**Table 5.**
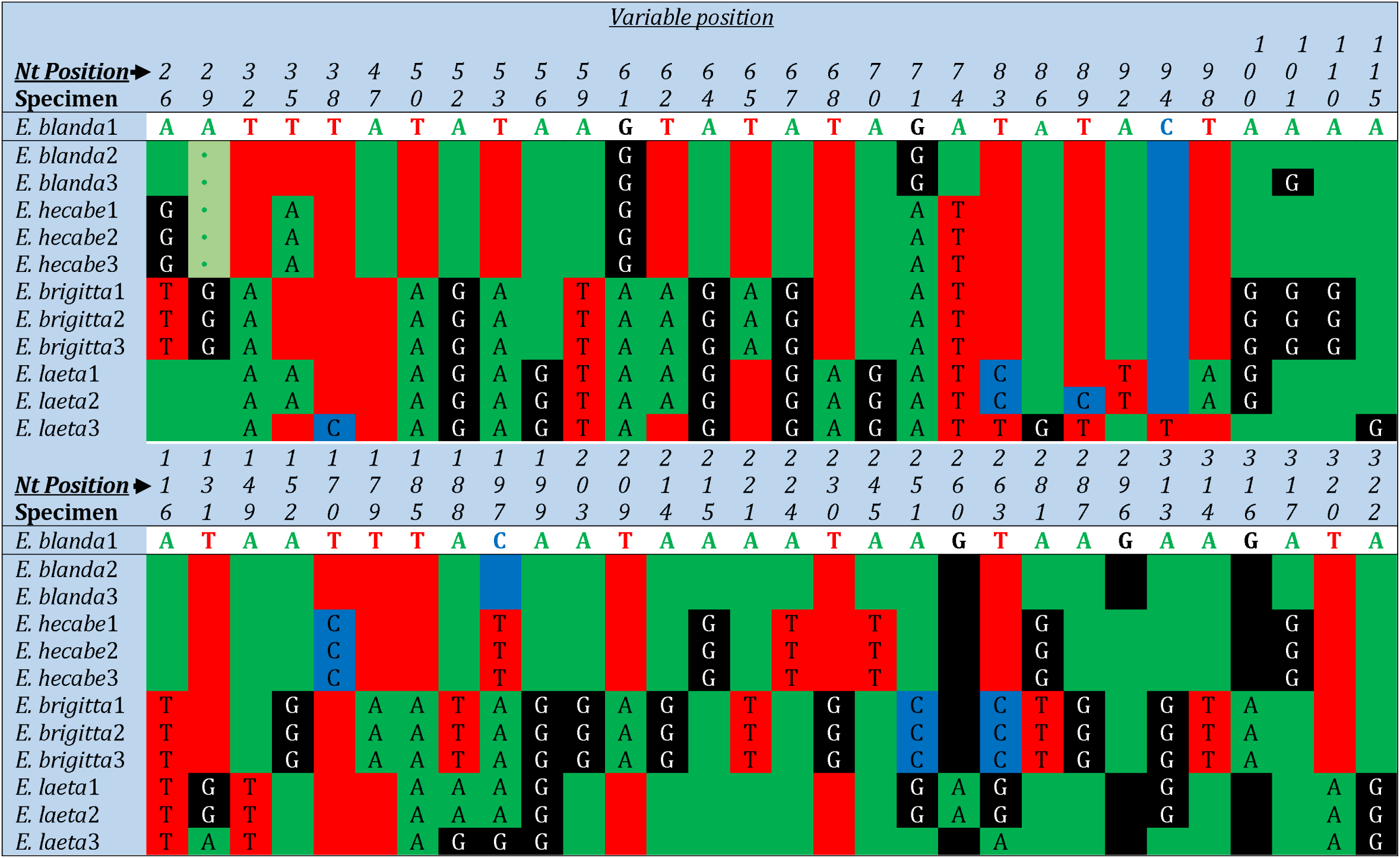
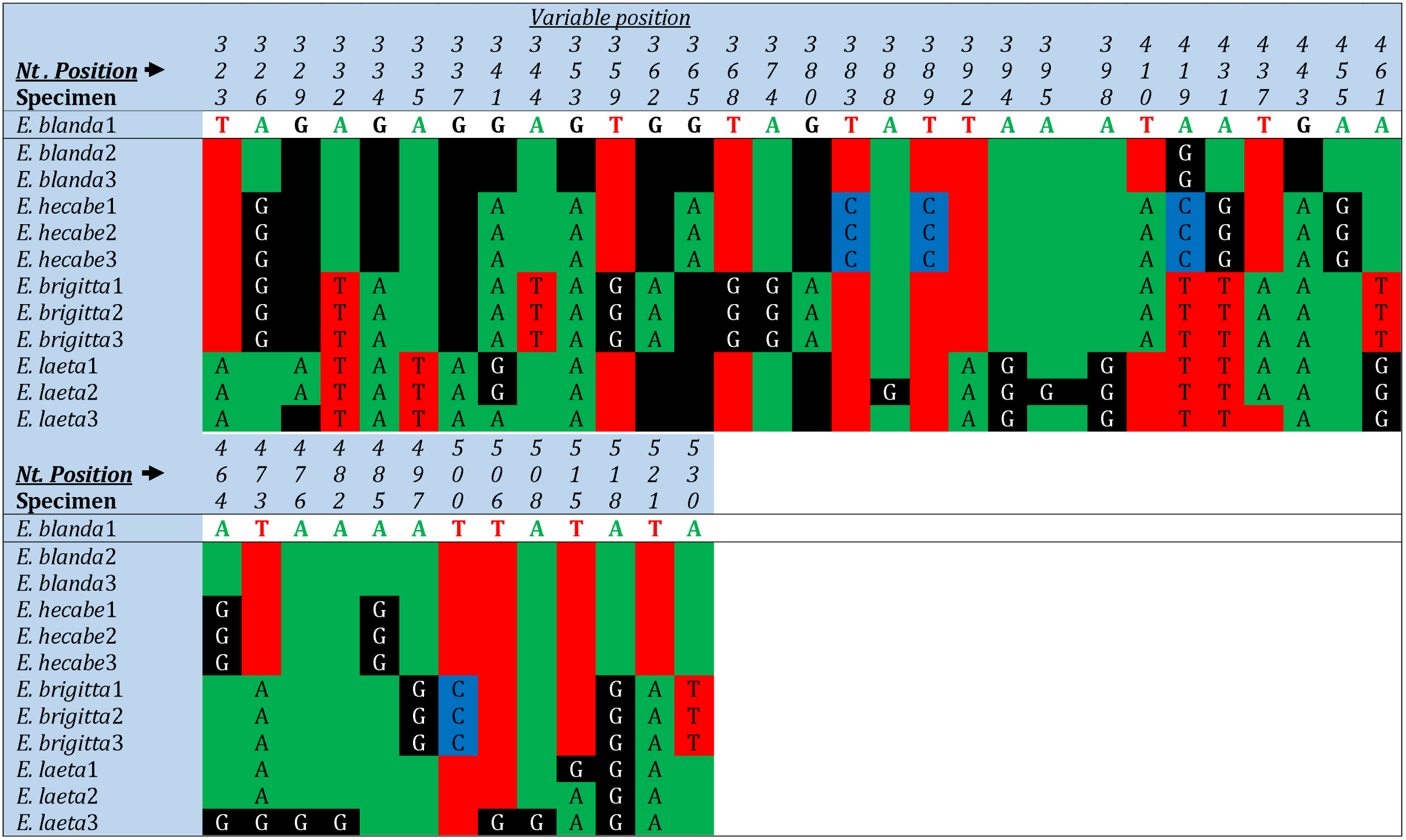
Observed COI sequnce based variable sites among four *Eurema* species present in Uttarakhand

The phylogenetic trees generated from COI sequences dataset by BI and ML approach tree were identical therefore; they merged into one tree depicting each tree individual support values. In tree topology, all twenty-one *Eurema* species recovered in two major clades (A and B) (Fig. 4). The clade A recovered as a monophyletic clade with strong bootstrap, although only seven of twenty-one species covered with two subclade (I and II) and was placed in the basal position as the sister to the rest of the clades of the *Eurema* species. In subclade I, only two species (*E. a. marginella* and *E. albula*) were present with strong bootstrap support (95%-100%), while subclade II covered five species (*E. nicippe, E. xantochlora, E. Mexicana, E. boisduvaliana* and *E. lisa*) with different sister clade although bootstrap support less (79%-84%).

**Figure 4:**
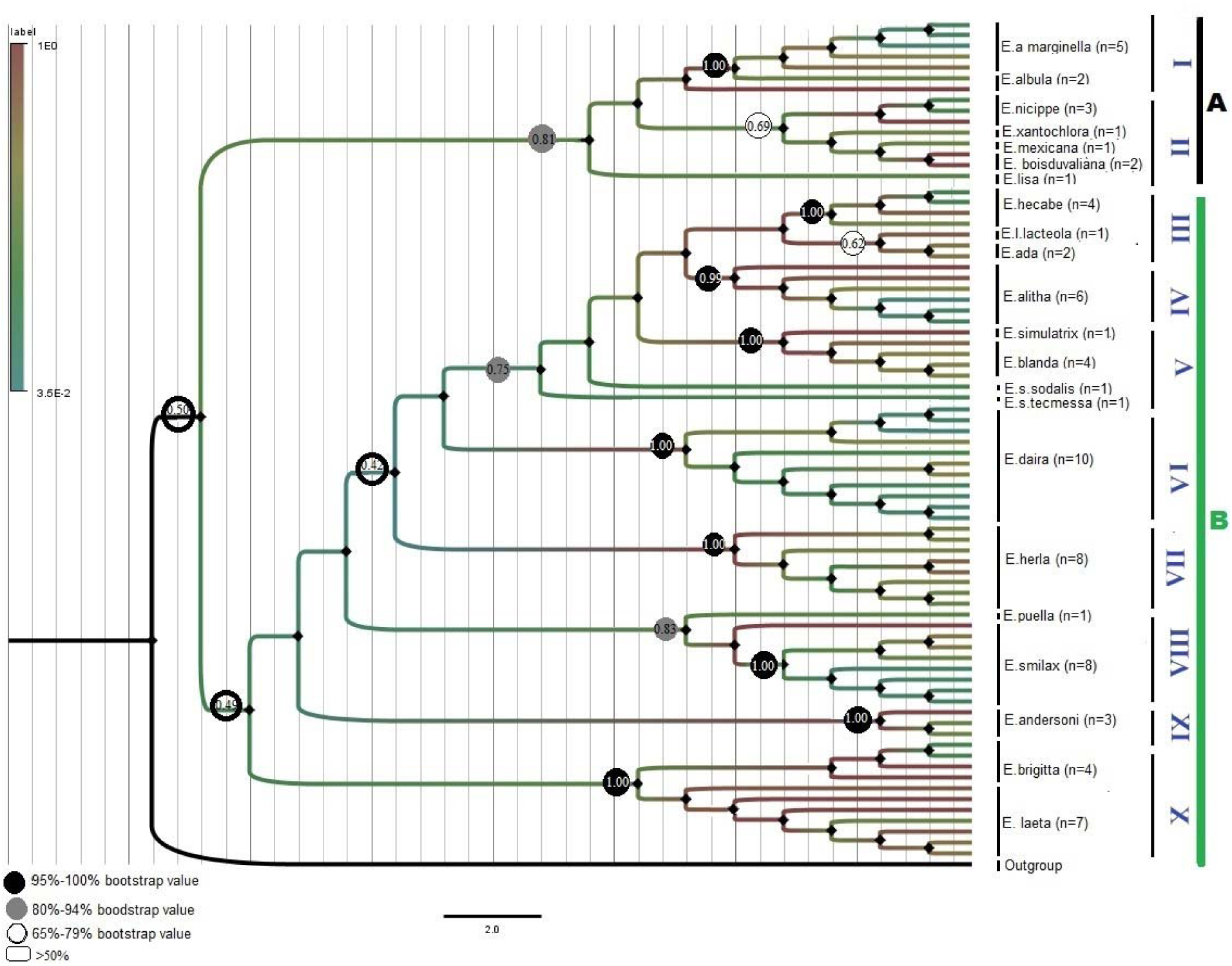
The ML and BI based phylogenetic tree showing the relationship between 21 species present under the genus *Eurema*. The ML tree reconstructed using the maximum likelihood (ML) method implemented in the PhyML program (v3.0) (Guindon, et al.,2010) and BI tree reconstruct using the MrBayes 3.1.231(Huelsenbeck JP & Ronquist F,2001).The circle with different colour showing the bootstrap value and number under circles represent BI clade posterior probability.

The clade B was a monophyletic clade and strongly supported bootstrap value, although fourteen *Eurema* species were also recovered into eight subclade (III-X) with supported bootstrap values. In subclade III, three of fourteen species (*E. hecabe, E. lecta* and *E.ada*) clustered with strong support value (95%-100%), although *E. lecta* and *E.ada* showed less support value (65%-79%), while in subclade V covered four species (*E. simulatrix, E. s. Sodalist, E. blanda* and *E. s. tecmessa*) with strong support value (95%-100%), though, two species (*E. simulatrix* and *E. blanda*) showed lesser support value (69%-75%). The subclade VIII and X consisted with four, two of each species i.e. (*E. puella* and *E. smilax*) and (*E. brigitta* and *E. leta*) with strongly supported bootstrap value (80%-94% and 95%-100%) respectively, while subclade IV, VI, VII and IX consisted with each species with high support value (95%-100%) respectively (Fig. 4).

Butterflies are affably the most attractive group of invertebrates and have been a source of inspiration for generations of natural historians and scientists. Subsequently, the classification of Lepidoptera (generic- and specific-level) is reasonably stable and the majority of taxa have named (Ackery et al. 1999). In the family Pieridae, *Eurema* with morphological synapomorphy is relatively unique and easy to be distinguished from the remaining genera. The five species (*E. hecabe, E. leata, E. brigitta, E. andersoni* and *E. blanda*) out of ca70 recognized species under the genus are distributed in Uttarakhand. In these DNA sequences of CO1 gene revealed that the obtained COI sequences are very helpful to discriminating the butterfly species of genus *Eurema*. The high interspecific and intraspecific variation between and within the Uttarakhand species support the unique DNA barcode of these species.

## CONCLUSION

The DNA sequences of COI gene revealed that the obtained COI sequences are very helpful to discriminating the butterfly species of genus *Eurema*. The present study samples correctly fall with the respective *Eurema* species without any ambiguities. Moreover, NJ clustering analysis of five Uttarakhand species showed well supported monophyletic clade of the sequences belonging to the same species without any overlap, even though these sequences are from the specimens separated by a large geographic distances (Fig 2, 3).

Some previous studies support long distance dispersal of butterflies (Craft et al. 2010, Gaikwad et al. 2011). Low sequence divergence and the sequences from the conspecifics butterflies of different biogeographically originated samples lies in monophyletic clades suggest long distance dispersal in Lepidoptera as well as in subjected genus in our study. This explains feasibility of DNA barcoding for species identification even when the species are separate by large geographic areas. Phylogenetic study of this genus, the two major clade (A and B) were confirmed, where the clade A represent the species of the Palearctic eco zone and the clade B represent species of the Indio-Malayan eco zone. Genus *Eurema* have large number of species (~70 species) and we included 21 species due to the lack of sequences, so there is need for further study of this genus to estimate the evolutionary pattern and deep phylogenetic relationship.

## ACKNOWLEDGMENT

The authors are grateful to Dr. K. Chandra, Director, Zoological Survey of India, Kolkata for their support and encouragement. We are very grateful to Mr. Pramod Kumar, Senior Zoologist, NRC, ZSI for helping in the initial identification of species. Authors also acknowledge the support provided by Officers and staff of NRC, ZSI, Dehradun. The authors also acknowledge the Department of Science and Technology (DST), Government of India, New Delhi for financial support under the Women Scientist Scheme (WOS-B) (DST/Disha/SoRF/022/2013).

